# The 1 –Cys peroxiredoxin, PRDX-6, suppresses a pro-survival response, including the Flavin monoxygenase, FMO-2, that protects against fungal and bacterial infection

**DOI:** 10.1101/2025.03.06.640586

**Authors:** Emma L. Button, Jake B. Lewis, Emilia Dwyer, Elise McDonald, Eloise Butler, Fiona Pearson, Elizabeth A. Veal

**Author notes:** These authors contributed equally to this manuscript (Jake B. Lewis, Emilia Dwyer). Correspondence: Dr Elizabeth Veal.

## Abstract

Reactive oxygen species (ROS)-induced cell damage contributes to many diseases. However, ROS also contribute to cell signaling and immune defences. As ubiquitous thiol peroxidases, peroxiredoxins (Prdx) play integral roles in balancing ROS functions. High levels of Prdx6 are associated with increased metastasis and resistance to chemotherapy, rendering Prdx6 a therapeutic target for treatment of a broad range of cancers. However, Prdx6, has additional activities, in lipid signalling and selenocysteine metabolism, and it remains unclear how Prdx6’s thiol peroxidase activity contributes to disease or ageing. Here we have investigated the role/s of Prdx6 in the stress responses and ageing of the nematode worm *Caenorhabditis. elegans*. Unexpectedly, we have found that *C. elegans* lacking *prdx-6,* have an increased resistance to oxidative stress and extended lifespan under some conditions. Moreover, *prdx-6* mutant worms are also more resistant to infection with two opportunistic human pathogens; the gram-positive bacteria *Staphylococcus aureus* and the dimorphic yeast *Candida albicans*. Our data suggest that increased ROS levels in *prdx-6* mutant worms lead to increased cell death in the germ line, and increased expression of the Flavin monooxygenase, FMO-2 in other tissues. FMO-2 has a conserved pro-survival function and is upregulated by the NHR-49(PPARα/HNF4) transcriptional regulator in response to various stresses, including peroxides and *S. aureus* infection. Here we reveal that *fmo-2* expression is also increased as an NHR-49-dependent protective response to *C. albicans*. Thus, in addition to its anti-ageing role, FMO-2 protects *C. elegans* against both fungal and bacterial infections. Accordingly, we propose that elevated *fmo-2* expression contributes to the increased stress resistance, lifespan and innate immunity of *prdx-6* mutant animals. These findings further illustrate the complex roles that ROS/PRDX can play in stress resistance, immunity and ageing.

**Highlights:**

- PRDX-6 is expressed in specific interneurons and intestine
- Loss of PRDX-6 increases germ cell apoptosis
- Loss of PRDX-6 increases *C. elegans* stress resistance, innate immunity and longevity
- Loss of PRDX-6 and infection increase expression of the flavin monooxygenase *fmo-2*
- FMO-2 protects against infection with the fungal pathogen *Candida albicans*

## Introduction

Reactive oxygen species (ROS) are generated as a by-product of aerobic metabolism and can cause profound damage to cellular macromolecules. Indeed, NADPH oxidases (Nox/Duox), ROS-generating systems, have evolved as an innate immune defence, exploiting ROS as powerful antimicrobials (For a review see (Sarr *et al,* 2018). Accordingly, enzymes and systems that reduce ROS levels play important roles in protecting against oxidative damage. Peroxiredoxins (Prdx) are amongst these ROS defences; reducing peroxides via reversible oxidation of a conserved, peroxide-reacting cysteine/s (Bolduc *et al*, 2021). Alterations in Prdx levels are linked with a number of diseases and ageing. For example, the mammalian 1-Cys peroxiredoxin (Prdx), Prdx6, has important roles in stress resistance (Eismann *et al*, 2009; Wang *et al*, 2003), apoptosis (Choi *et al*, 2011; Kim *et al*, 2011) and is up-regulated in many cancers (Karihtala *et al*, 2003; Lehtonen *et al*, 2004; Hu *et al*, 2020). Moreover, high levels of Prdx6 expression are associated with increased metastasis and resistance to chemotherapy making them candidate chemotherapeutic targets to treat a range of cancers (Chen *et al*, 2021; Ho *et al*, 2010; Huang *et al*, 2018; Jo *et al*, 2013; Pak *et al*, 2011; Yun *et al*, 2015; Sahu *et al*, 2016; Hu *et al,* 2020).

However, in addition to their peroxidase activity, eukaryotic Prdx have other activities that may contribute to their *in vivo* function. For example, 2-Cys Prdx directly and indirectly mediate some of the positive signaling functions of low levels of peroxide (For reviews see (Bolduc *et al*., 2021; Netto & Antunes, 2016). Moreover, the mammalian 1-Cys Prdx, Prdx6, has additional signaling activities: a Ca^2+^ independent phospholipase A_2_ (iPLA_2_) activity and a lysophosphatidylcholine acyl transferase (LPCAT) activities (for a review see (Fisher, 2018). The peroxidase, iPLA_2_ and LPCAT activities have distinct active sites, with the peroxidase activity centered around a catalytic cysteine and the iPLA_2_ activity centered around a catalytic serine and a lipase motif. Studies in mice have suggested that peroxidase, lipase and acylation activities all contribute to Prdx6’s physiological functions and roles in disease (Ho *et al*., 2010; Li *et al*, 2015; Lien *et al*, 2012). It has been proposed that the lipid-binding capacity of Prdx6 allows it to protect cell membranes against ROS-induced damage by detoxifying phospholipid hydroperoxides (Fisher *et al*, 2018). However, Prdx6 was recently shown to protect against lipid peroxides indirectly; by facilitating the incorporation of selenium into selenoproteins, such as the glutathione peroxidase, Gpx4 (Fujita *et al*, 2024; Chen *et al*, 2024; Ito *et al*, 2024).

Indeed, the presence of multiple peroxiredoxins and 30 selenoproteins, including 4 glutathione peroxidases make the contributions of Prdx6’s different activities to ageing and disease, more challenging to dissect in mammals. In contrast, the genome of the nematode worm *Caenorhabditis elegans* encodes only 3 peroxiredoxin genes (**Fig. 1A**) and only a single selenoprotein, the thioredoxin reductase, TRXR-1, which is redundant with glutathione reductase (Stenvall *et al*, 2011). Moreover, *C. elegans* is also a highly useful system to investigate the roles of ROS in signalling, ageing and infection (for a review see (Miranda-Vizuete & Veal, 2017). For example, the *C. elegans* orthologue of the mammalian Nrf2 transcription factor, SKN-1, plays an important role in mediating many of the positive effects of ROS (For a review see (Blackwell *et al*, 2015)). In addition, the NHR-49 transcription factor, that controls similar genes to mammalian PPARα, also responds to ROS, activating a protective transcriptional response to peroxide (Goh *et al*, 2018).

**Fig. 1.**
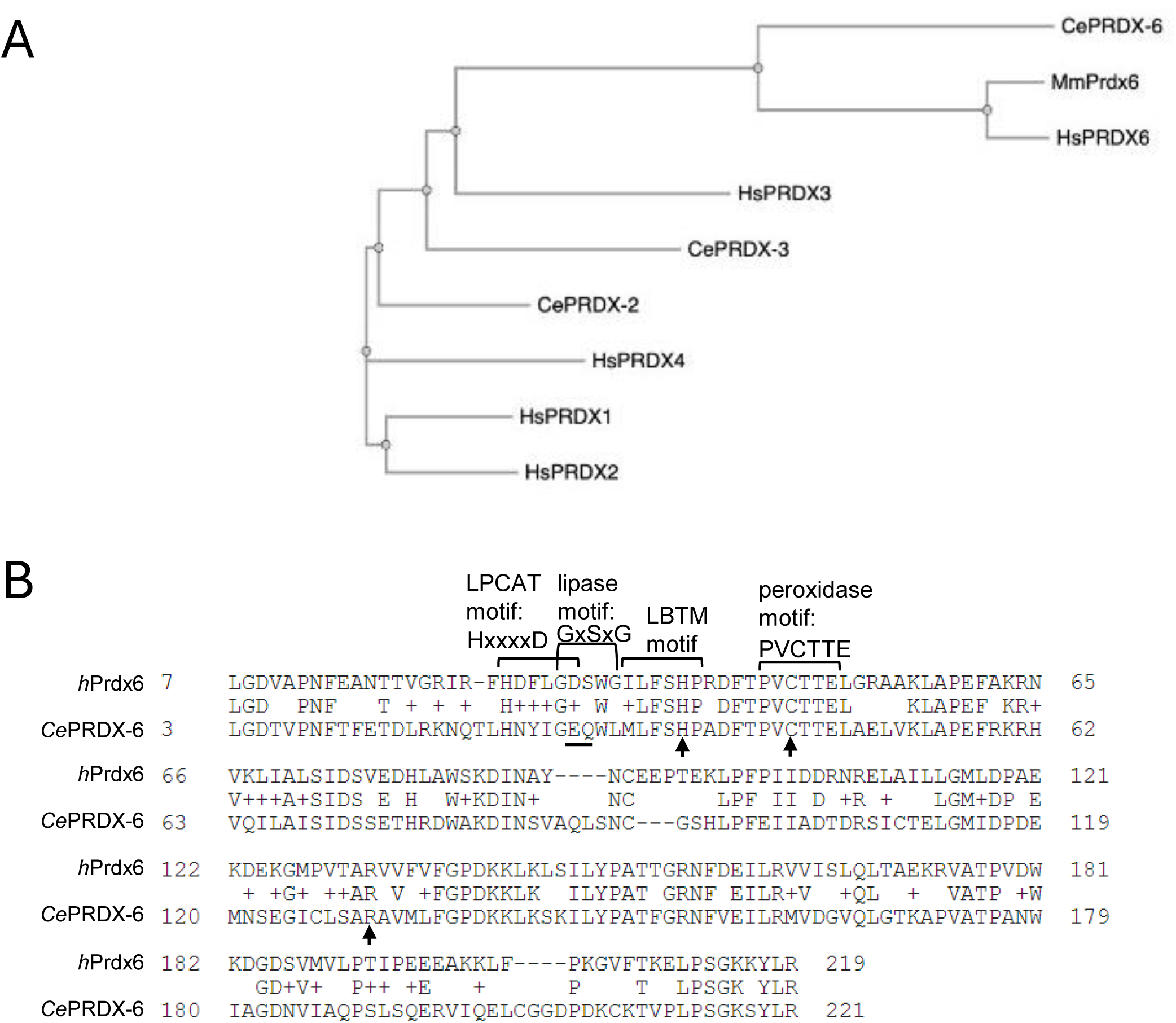
Sequence comparisons reveal that *C. elegans* PRDX-6 is [A] more closely related to mammalian Prdx6 proteins than other 1 and 2-Cys Prdx and [B] contains the peroxidase motif present in Prdx6 family but lacks the phospholipase A2 or LPCAT motifs found in human Prdx6. **[A]** Phylogenetic tree generated using ClustalW2 software demonstrating the relationships between the *C. elegans* and mammalian 1-cys peroxiredoxin, CePRDX-6, the human PRDX-6 (HsPRDX-6) and the *Mus musculus* PRDX-6 (MmPRDX-6), and the 2-cys peroxiredoxins, CePRDX-2 and CePRDX-3 and HsPRDX1, HsPRDX2, HsPRDX3 and HsPRDX4. **[B]** Alignment of CePRDX-6 and human Prdx6 amino acid sequences indicates that the peroxidase motif, PVCTTE, that is unique to Prdx6 family, and the catalytic triad C47-H39 - R132 (arrows) are both present in human and *C. elegans* PRDX-6. The LPCAT motif, HxxxxD and the lipase motif, GxSxG, required for acylation of lysophosphatidylcholine and for Ca^2+^ independent phospholipase A2 (iPLA2) activity of mammalian Prdx6 respectively, are absent from *C. elegans* PRDX-6, with the critical aspartate for LPCAT activity in the LPCAT motif replaced with glutamate and the critical serine for phospholipase activity in the lipase motif replaced with a glutamine (underlined) in *C. elegans* PRDX-6.

Previous studies have suggested that the mitochondrial 2-Cys Prdx, PRDX-3, does not play an important role in protecting against oxidative damage, and *prdx-3* mutant animals have a wild-type lifespan (Olahova et al., 2008; Ranjan *et al*, 2013). In contrast, the single cytosolic 2-Cys Prdx, PRDX-2, plays several important physiological roles (Bhatla & Horvitz, 2015; Olahova *et al*, 2008; Olahova & Veal, 2015). This includes tissue-specific roles, with intestinal PRDX-2 protecting against the acute toxicity of hydrogen peroxide, whereas PRDX-2 is required in specific neurones for hydrogen peroxide-induced inhibition of feeding (Bhatla & Horvitz, 2015). Consistent with a positive role in promoting ROS signal transduction, PRDX-2 is also required for arsenite and metformin-induced activation of the p38 MAPK, PMK-1 (De Haes *et al*, 2014; Olahova *et al*., 2008). However, PRDX-2 limits resistance to some oxidative stress conditions, including arsenite, by promoting insulin secretion that inhibits the activity of SKN-1(Nrf2) and DAF-16(FOXO) (Olahova *et al*., 2008; Olahova & Veal, 2015). Notably, despite reduced insulin signalling and increased SKN-1 and DAF-16 activity, *prdx-2* mutant animals are short-lived (Olahova *et al*., 2008; Olahova & Veal, 2015). However, it is unclear whether the *C. elegans* 1-Cys Prdx, PRDX-6 plays important roles under physiological or stress conditions (Isermann *et al*, 2004). The goal of this study was to address this question.

Comparison of the *C. elegans* PRDX-6 sequence with the human Prdx6 sequence demonstrates that *C. elegans* PRDX-6 contains the peroxidase domain and histidine important for lipid binding, but the catalytic serine required for iPLA_2_ activity is not conserved (**Fig. 1B**). Moreover, *C. elegans* only incorporate selenocysteine into a single protein, the thioredoxin reductase TRXR-1, which is redundant with glutathione reductase (Stenvall *et al*, 2011). This provided us with the opportunity to investigate the contribution that the peroxidase activity plays to the *in vivo* biological role/s of a 1-Cys Prdx in a whole animal model. Consistent with PRDX-6’s peroxidase activity playing an important role in removing endogenous peroxides we find that loss of *prdx-6* causes detectable increases in intracellular levels of ROS and germ cell apoptosis. However, unexpectedly, loss of PRDX-6 increases the oxidative stress resistance, lifespan and innate immunity of *C. elegans*. Our data suggest that the increased resistance of *prdx-6* mutant *C. elegans* to oxidative stress and pathogenic infection may be due to increased ROS levels leading to increased expression of peroxide-induced genes, including the pro-longevity *fmo-2* gene (Leiser *et al*, 2015). FMO-2 is also induced as an NHR-49-dependent response to bacterial infection (Dasgupta *et al*, 2020; Wani *et al*, 2021). Here we reveal that *fmo-2* also protects *C. elegans* against the fungal pathogen *Candida albicans.* Thus, we propose that loss of PRDX-6 increases activity of NHR-49(HNF4/PPARα) to prime *C. elegans* innate immune defences against opportunistic human pathogens.

## Results

### PRDX-6 is expressed in the intestine, I2 and I4 interneurones

To determine where PRDX-6 was expressed, transgenic animals were generated in which the 2.5kb promoter region upstream of the *prdx-6* orf was used to drive the expression of the first 22 amino acids of PRDX-6 in frame with GFP (*prdx-6_prom_::gfp*) (**Fig. 2A**). Consistent with high throughput single cell mRNA expression analyses (Taylor *et al*, 2021), *prdx-6_prom_*::GFP expression was detected in the intestine and specific neurones within the head in all larval stages and adults and the gonadal sheath of adult hermaphrodites (**Fig. 2B**). The intestinal expression of *prdx-6_prom_*::GFP, was greatest in the most anterior and posterior intestine, consistent with the expression of many stress-induced genes (An & Blackwell, 2003; Hattori *et al*, 2013). In the head *prdx-6_prom_*::GFP was expressed specifically in three neurones. Two of the cell bodies were in the anterior bulb with symmetrical axonal projections, indicating they represented a pair of neurones (**Fig. 2C**). To determine the identity of these neurones, we compared the expression of *prdx-6_prom_::gfp* with that of the tryptophan hydroxylase, TPH-1, that is expressed in serotonergic neurones, including the neurosecretory-motor neurones (NSM) neurones. Co-expression of *prdx-6_prom_*::GFP and *tph-1*::dsRed2, suggested that *prdx-6_prom_*::GFP is expressed in the I2L and I2R inter-neurones, that are adjacent to the NSM neurones in which TPH-1 is expressed, as well as a third inter-neuron with a cell body in the terminal bulb and axons projecting to the anterior bulb that is likely to be I4 (**Fig. 2C-D**).

**Fig. 2.**
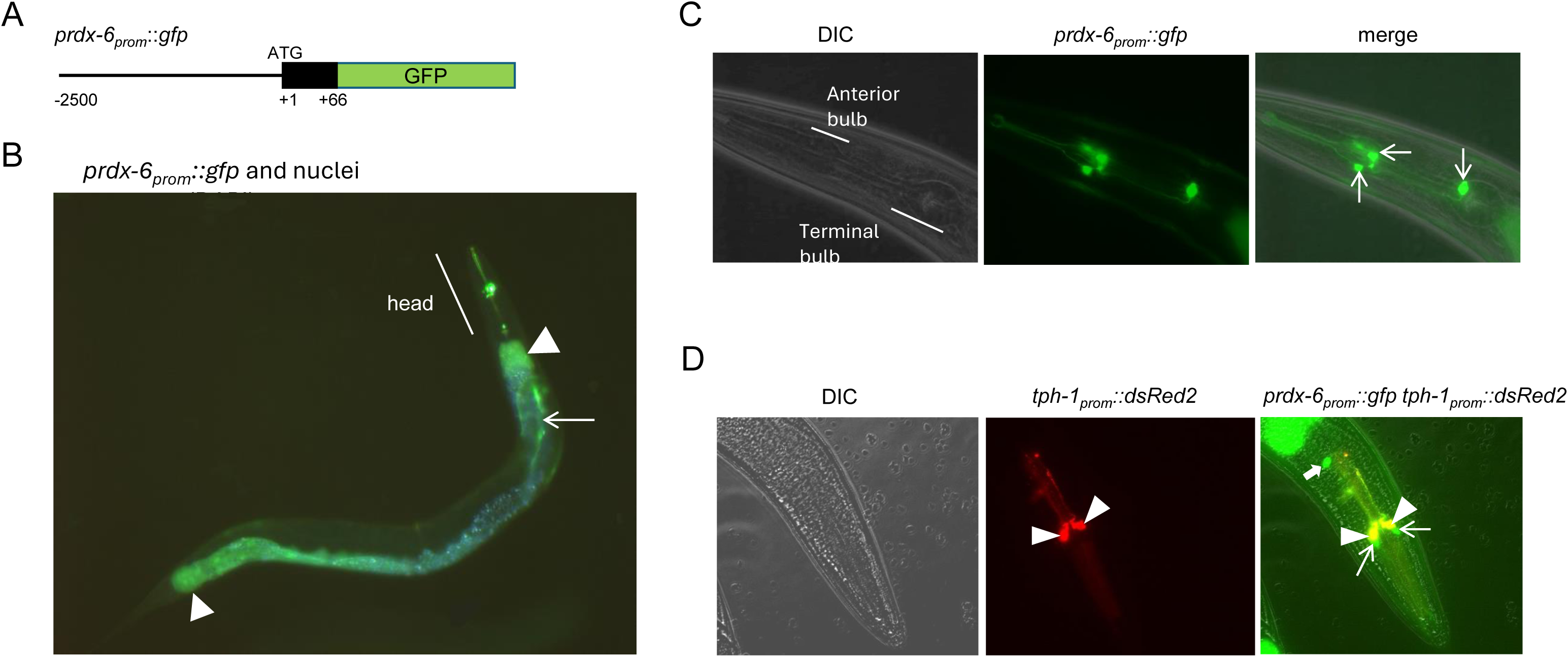
Analysis of transgenic animals expressing a transcriptional reporter reveals that PRDX-6 is expressed in the intestine, gonadal sheath, I2 and I4 neurones. **[A]** Schematic representation of the *prdx-6P::gfp* transcriptional reporter **[B]** Analysis of *prdx-6P::gfp* expression in a young adult worm, nuclei were imaged by staining with DAPI (blue) and a merged images of green and blue channels is shown. Arrowheads indicate ends of intestine, arrow indicates gonadal sheath. **[C]** Analysis of *prdx-6P::gfp* expression in head. GFP-positive neuronal cell bodies are indicated by arrows on merged image of DIC and GFP. **[D]** Analysis of *prdx-6P::gfp* expression in head of *C. elegans* co-expressing *tph-1P*::dsRed2 in the NSM cell bodies (labelled with triangles). Thin arrows on merged image (right hand panel) indicate *prdx-6*p::GFP-positive cell bodies adjacent to the NSM (interneurones I2L and I2R). DsRed2 also fluoresces under the GFP excitation wavelength therefore *tph-1*p::DsRed2-positive NSM neurones appear orange on the merged image. A thick arrow indicates the third, unidentified, *prdx-6*p::GFP-positive neuronal cell body.

### Loss of prdx-6 increases resistance to oxidative stress and lifespan at 15°C

To analyse the function of PRDX-6, an RNAi clone was generated targeting PRDX-6. Western blot analysis, using anti-PRDX-6 antibodies, detected a band, consistent with the expected size of PRDX-6 (25.6kD), in lysates from control animals that was absent from *prdx-6* RNAi-treated animals, confirming that *prdx-6* RNAi was effective at reducing PRDX-6 levels (**Fig. S1A**). Two *prdx-6* mutant alleles were also obtained from the National Bioresource Project (NBRP), *prdx-6* (*tm4225*) and *prdx-6* (*tm4284*) containing a 442bp delete and a 4bp insertion in the second intron and a 356bp deletion in the third exon respectively (**Fig. S1B**). Analysis of protein lysates from wild type, and outcrossed *prdx-6* (*tm4225*) and *prdx-6* (*tm4284*) mutant worms using anti-PRDX-6 antibodies confirmed the absence of PRDX-6 from either *prdx-6* (*tm4225*) or *prdx-6* (*tm4284*) mutant worms suggesting that both *prdx-6* (*tm4225*) and *prdx-6* (*tm4284*) are null alleles (**Fig. S1C**).

Although I2 neurones are important for the hydrogen peroxide-induced inhibition of feeding, we found no evidence that PRDX-6 was important for this behavioural response to peroxide (**Fig. S2**)(Bhatla &Horvitz, 2015). However, consistent with the peroxidase activity of PRDX-6 acting as a barrier to endogenous ROS, we observed increased DCFDA staining in *prdx-6* mutant animals suggesting ROS levels were increased (**Fig. 3A**). Peroxide treatment increases the number of apoptotic corpses detected in the *C. elegans* germline. Notably, the increased number of apoptotic corpses in *prdx-6* mutant germlines compared with wild type animals, suggesting that PRDX-6 protects against ROS-induced apoptosis (**Fig. 3B-C**) (Fausett *et al*, 2021). However, when we examined the survival of wild type, *prdx-6* (*tm4225*) and *prdx-6* (*tm4284*) mutant worms on plates containing toxic levels of hydrogen peroxide and arsenite, we found that *prdx-6* mutant animals were significantly more resistant to treatment with both oxidative stress agents (**Fig. 4A-B**).

**Figure 3.**
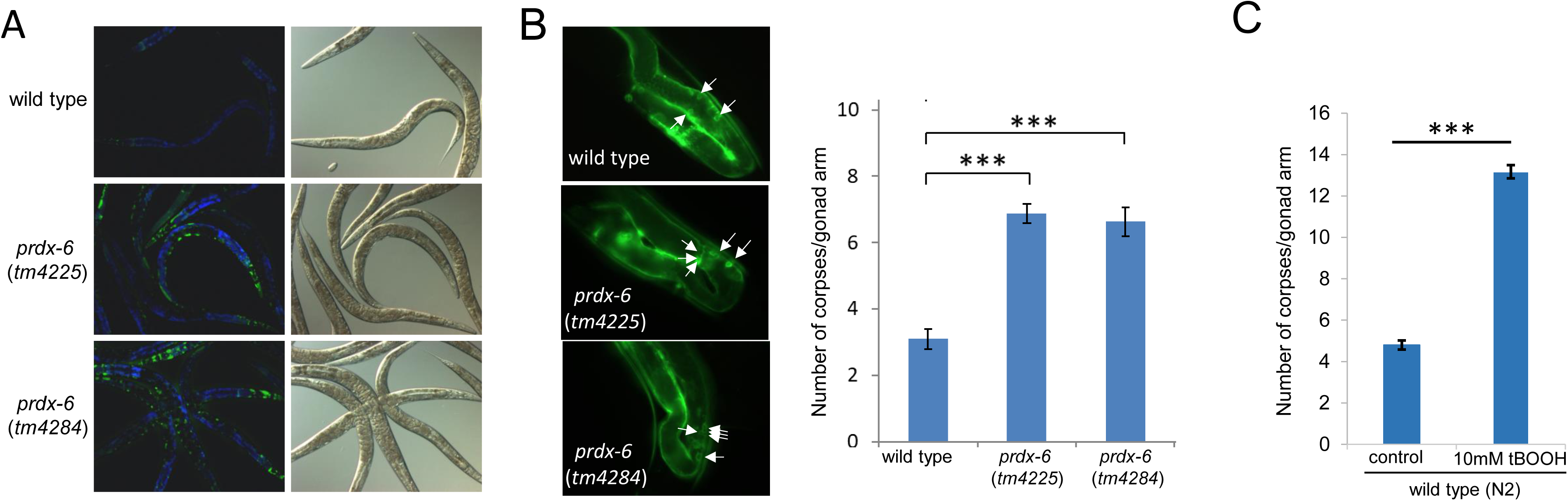
Levels of ROS and apoptosis are increased in *prdx-6* mutant *C. elegans*. **[A]** wild type, *prdx-6* (*tm4225*) and *prdx-6* (*tm4284*) mutant worms were stained with the ROS probe H_2_DCFDA and the fluorescence observed using fluorescence spectroscopy. Over 30 worms were observed in at least three independent experiments and representative DIC and fluorescent images are shown, with DCF fluorescence in green and autofluorescence in blue. **[B]** Wild type(N2) *ced-1*::GFP, *prdx-6* (*tm4225*) *ced-1*::GFP and *prdx-6* (*tm4284*) *ced-1*::GFP mutant worms were used to visualise engulfed apoptotic corpses in the germline (indicated by arrows). The number of apoptotic corpses in one gonad arm were counted in young adult wild type (N2) *ced-1*::GFP, *prdx-6* (*tm4225*) *ced-1*::GFP and *prdx-6* (*tm4284*) *ced-1*::GFP mutant animals or **[C]** before or following 90 min exposure (followed by 60 min recovery) of wild type(N2) *ced-1*::GFP to 10mM tert-butyl hydroperoxide (tBOOH) at 24°C. Mean values are shown from groups of 25-30 animals. P values were determined using Students T-test. *** p<0.001.

**Fig. 4.**
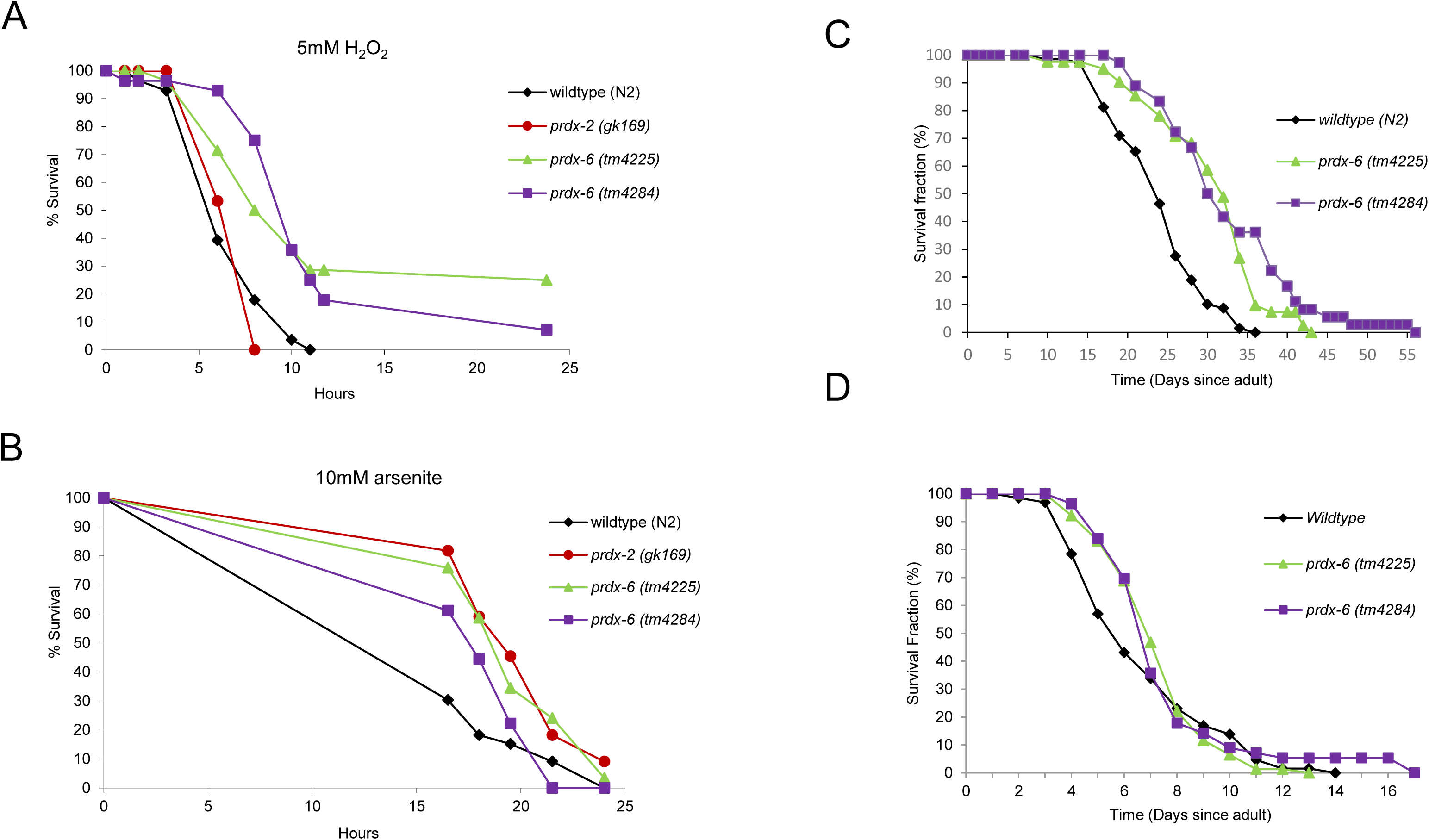
Loss of PRDX-6 increases survival under acute oxidative stress conditions and lifespan at 15°C but not 25°C. The survival of L4 larval stage wild type, *prdx-2 (gk169*), *prdx-6* (*tm4225*) and *prdx-6* (*tm4284*) mutant animals was monitored at the indicated time points [A-B] on plates containing either **[A]** 5.0mM H_2_O_2_ or **[B]** 10.0mM arsenite. These experiments were carried out three times with similar results and a representative experiment is shown. n=23-35 in each group. Log rank analysis indicates that the survival of *prdx-6* mutant animals is significantly increased compared with wild-type (p<0.002). **[C-D]** The survival of wild type (N2) (n=69*), prdx-6* (*tm4225*) (n=41) and *prdx-6* (*tm4284*) (n=36) mutant worms was monitored on live *E. coli* OP50 at **[C]** 15°C or **[D]** 25°C. Experiments were repeated 3 times and representative experiments are shown. Mean and median survival of wild type (N2), *prdx-6* (*tm4225*) and *prdx-6* (*tm4284*) mutant worms and log-rank analysis of *prdx-6* (*tm4225*) and *prdx-6* (*tm4284*) mutant worms compared with wild type indicated that differences in survival were statistically significant at 15°C but not 25°C (**Table S1-3**).

Our discovery that *prdx-6* mutant worms have an increased resistance to H_2_O_2_ contrasts sharply with the increased H_2_O_2_ sensitivity of animals lacking PRDX-2 (Oláhová *et al*, 2008). Moreover, whereas loss of *prdx-2* reduces the lifespan of *C. elegans* at 15°C (Oláhová *et al*., 2008) *prdx-6* mutant animals were long-lived at 15°C **(Fig. 4C, Table S1)**. The effect of PRDX-2 on lifespan is temperature-dependent, with wild-type and *prdx-2* mutant worms having similar lifespans at 25°C (Oláhová *et al*., 2008; Isermann et al 2004). Intriguingly, loss of PRDX-6 did not significantly alter the lifespan of animals maintained at 25°C either, suggesting that like PRDX-2, PRDX-6, has a temperature-dependent role in longevity (**Fig. 4D, Table S2-3**). Overexpression of PRDX-2 increases the resistance of wild-type animals to both arsenite and H_2_O_2_ (Oláhová *et al*., 2008). This raised the possibility that a compensatory increase in PRDX-2 levels might be responsible for the increased lifespan and oxidative stress resistance of *prdx-6* mutant *C. elegans*. However, PRDX-2 levels were similar in wild type, *prdx-6* (*tm4225*) and *prdx-6* (*tm4284*) mutant worms, both before and after exposure to H_2_O_2_ (**Fig. S3**).

### Loss of PRDX-6 increases the resistance of *C. elegans* to *Staphylococcus aureus* and *C. albicans* infection but does not protect against Salmonella Typhimurium

There were a number of possible explanations for the temperature-dependent effect of PRDX-6 on lifespan. For instance, a major lifespan-limiting factor at 25°C is the increased growth and pathogenicity of the *E. coli* food source at these higher temperatures (Oláhová *et al*., 2008). Moreover, while ROS play an important role in protecting *C. elegans* against some pathogens (Liu *et al*, 2019; Tiller & Garsin, 2014) ROS generated during infection, have been proposed to be partially responsible for the premature death of *C. elegans* infected with the gram negative bacteria, S. Typhimurium (Aballay *et al*, 2000; Labrousse *et al*, 2000). Indeed, the survival of *prdx-6* mutants exposed to *S*. Typhimurium, which establishes a persistent infection in the intestine that causes the premature death of the worm, was reduced compared with wild-type animals (**Fig. 5A, Table S4**). However, in contrast, loss of PRDX-6 significantly increased the resistance of *C. elegans* to infection with two other human pathogens, the gram positive bacteria, *Staphylococcus aureus* and the dimorphic fungus *C. albicans* (**Fig. 5B-C, Tables S5-6**). Intriguingly, this suggests that PRDX-6 has pathogen-specific roles in innate immunity. The increased resistance of *prdx-6* mutant *C. elegans* to infection with *S. aureus* and *C. albicans*, both of which kill *C. elegans* more quickly than S. Typhimurium, was consistent with the increased resistance of these animals to the acute toxicity of arsenite and peroxide (**Fig. 4A-B**). This raised the possibility that loss of PRDX-6 activates a stress-protective response that increases the resistance to a variety of acute stresses, but does not protect against a more chronic infection with S. Typhimurium.

**Fig. 5.**
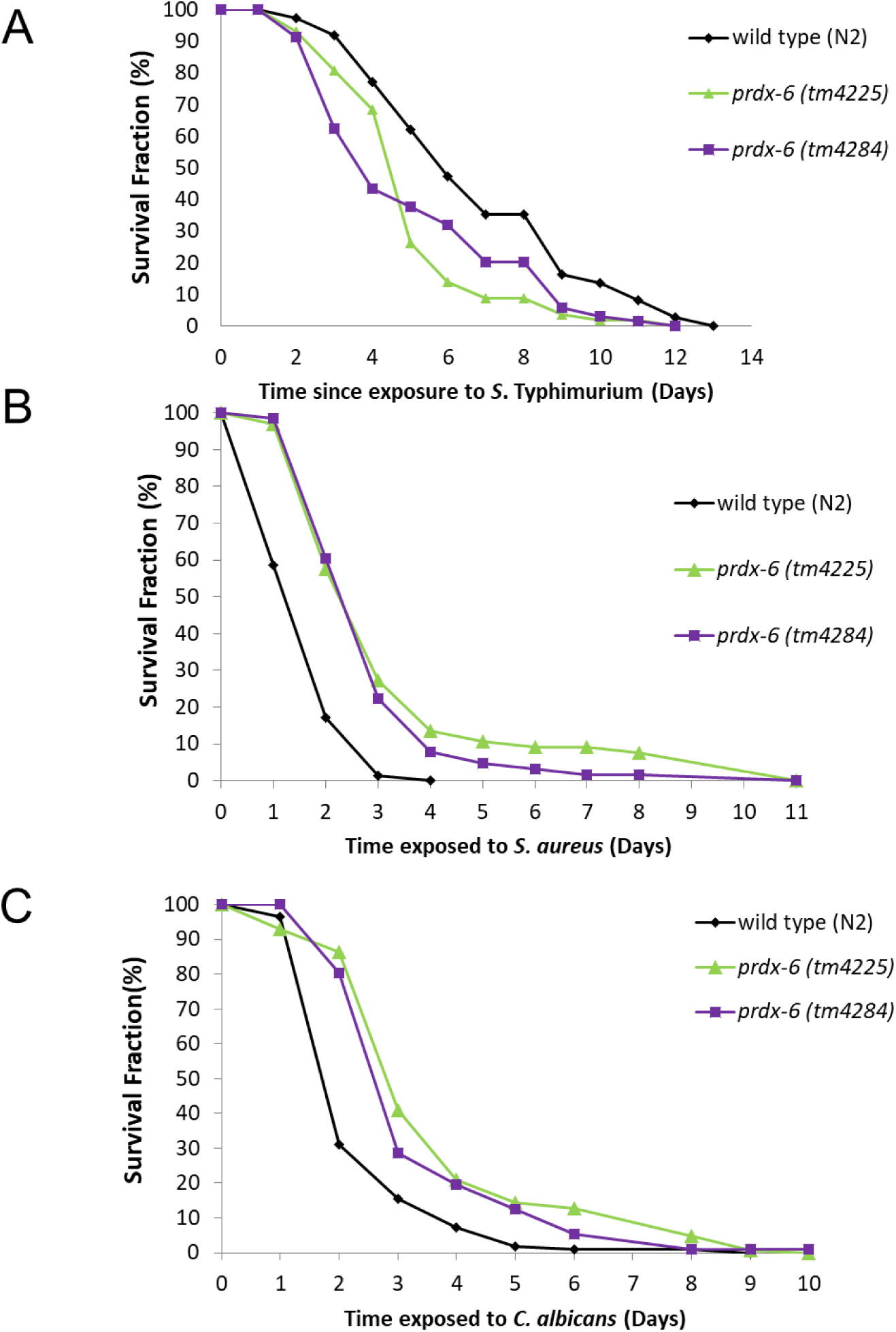
Loss of *prdx-6* increases resistance to infection by *S. aureus* and *C. albicans* but not *Salmonella Typhimurum*. The survival of young adult wild type (N2), *prdx-6* (*tm4225*) and *prdx-6*(*tm4284*) mutant animals transferred on to plates seeded with **[A]** *S*. Typhimurium (SL1344) **[B]** *S. aureus* (NCTC 8325) or **[C]** *C. albicans* (SN148). These experiments were carried out with >50 animals per group and repeated three times with similar results. Representative experiments are shown. The mean survival of wild type, *prdx-6* (*tm4225*), *prdx-6* (*tm4284*) and P-values were determined by log-rank analysis (**Tables S4-6**).

### Loss of PRDX-6 and exposure to *C. albicans* and *S. aureus* increase expression of the Flavin monoxygenase, FMO-2 which protects against both pathogens

Next, we investigated the basis for the increased resistance of *prdx-6* mutant *C. elegans* to infection with *S. aureus* or *C. albicans*. Arsenite, peroxide and infection are amongst the stresses that have been shown to activate the *C. elegans* Cap n Collar transcription factor SKN-1, that mediates protective transcriptional responses. Indeed, increased SKN-1 activity is important for resistance to arsenite, protects against pathogens and can extend lifespan (Blackwell *et al*., 2015). However, we could not find any evidence that the increased ROS in animals lacking *prdx-6* was sufficient to increase expression of a SKN-1-dependent reporter gene, expressing GFP from the *gcs-1* promoter (**Fig. S4**). Notably, one of the most highly peroxide-induced genes the Flavin monoxygenase, *fmo-2*, is activated in a *skn-1*-independent manner by the nuclear hormone receptor NHR-49 (Goh *et al*., 2018). Although wild-type and *prdx-6* mutant *C. elegans* appeared to contain similar nuclear levels of NHR-49 (**Fig. S5**), it was plausible that loss of PRDX-6 might increase NHR-49 activity by increasing the availability of an oxidised lipid ligand (Doering *et al*., 2023). Hence, we tested whether the increased ROS in *prdx-6* mutants might be sufficient to increase the expression of *fmo-2*. Notably, we found that, although the levels of expression of an *fmo-2P::gfp* reporter are normally low, intestinal levels of expression were significantly increased in animals lacking *prdx-6* (**Fig. 6A**). Increased FMO-2 expression increases the lifespan of *C. elegans* maintained on non-proliferating *E. coli*, eliminating this reflecting an anti-microbial effect (Leiser *et al*., 2015). However, our finding suggested that FMO-2 might also protect against infection, and that increased expression of *fmo-2* might contribute to the increased resistance of *prdx-6* mutant *C. elegans* to infection with *S. aureus* or *C. albicans*.

**Fig. 6.**
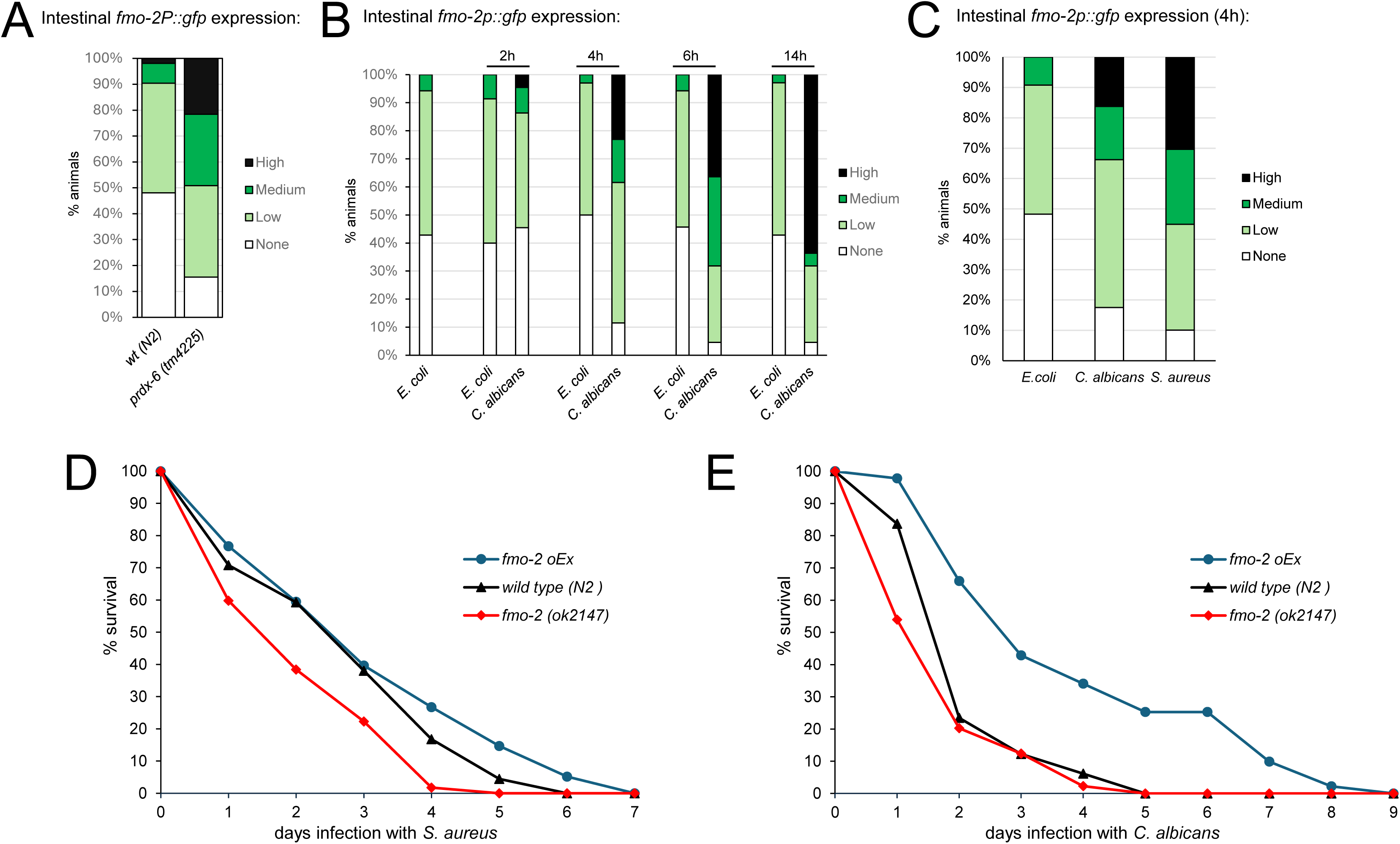
FMO-2 expression is induced by loss of *prdx-6* and protects against infection with *C. albicans* and *S. aureus*. The intestinal expression of a *fmo-2P::gfp* reporter gene was **[A]** increased in *prdx-6* (*tm4225*) mutant compared with wild-type animals (Chi^2^ test p= 8.89889E-07) **[B-C]** rapidly increases following transfer of *C. elegans* to **[B]** *C. albicans* or **[C]** *S. aureus* compared with transfer to *E. coli* control. **[D-E]** The survival of wild-type (N2), *fmo-2* (*ok2147*) mutant and *fmo-2* over-expressing (*fmo-2 OEx*) C*. elegans* following exposure to **[D]** *S. aureus* or **[E]** *C. albicans.* The experiments shown are representative of at multiple (>3) independent experiments involving approximately 80-150 animals in each group. The mean survival of wild type, *fmo-2* (*ok2147*) and *fmo-2oEx* and P-values were determined by log-rank analysis (**Tables S7-8**).

*C. albicans* and *S. aureus* infection initiate both distinct and overlapping *C. elegans* transcriptional responses (Pukkila-Worley *et al*, 2011). Consistent with the hypothesis that increased expression of *fmo-2* might be an important innate response to infection, *fmo-2* is very highly induced in response to *Staphylococcus aureus* (Irazoqui *et al*, 2010; Wani *et al*., 2021). Moreover, although *fmo-2* was not amongst the genes found to be induced in response to *C. albicans* in previous studies (Pukkila-Worley *et al*., 2011), exposure of wild-type *C. elegans* to *C. albicans* also caused a rapid increase in the intestinal expression of *fmo-2::gfp* (**Fig. 6B**). This increase preceded any visible evidence of *C. albicans* infection but was not apparent when worms were exposed to heat-killed *C. albicans*, suggesting that induction was due to a non-denaturable signal emanating from the yeast (**Fig. S6**). Indeed, we observed a similar increase in intestinal *fmo-2p::gfp* expression when animals were exposed to either *C. albicans* or *S. aureus* indicating that *fmo-2* induction is part of a common response to these 2 pathogens (**Fig. 6C**). As previously reported, the ability of *fmo-2* mutant animals to survive infection with *S. aureus* was significantly compromised (**Fig. 6D, Table S7)** (Wani *et al*., 2021). There was also a small, but consistent, increase in the susceptibility of *fmo-2* mutant animals to *C. albicans*, particularly in the early stages of infection (**Fig. 6E, Table S8**). Conversely, the survival of animals expressing multiple copies of the *fmo-2* gene infected with *C. albicans* was significantly prolonged compared with wild-type (p<0.001) (**Fig. 6E**) and there was a similar correlation between FMO-2 levels and survival of *S. aureus*-infected animals (**Fig. 6D**). This suggests that this flavin monoxygenase, is part of a common, protective innate immune response to fungal and bacterial pathogens.

The NHR-49 transcriptional regulator is activated by tBOOH and required for increased *fmo-2* expression in response to peroxide, fasting and *S. aureus* infection (Goh *et al*., 2018; Wani *et al*., 2021). Similarly, *C. albicans*-induced increases in *fmo-2p*::GFP expression were ablated in *nhr-49* mutant *C. elegans* (**Fig. 7A**). Accordingly, we propose that the increased ROS present in *prdx-6* mutant animals may increase NHR-49 activity driving the expression of *fmo-2* and other genes (Oliviera *et al*., 2009; Goh *et al*., 2018; Wani *et al*., 2021) responsible for their increased survival (**Fig. 7B**).

**Fig. 7.**
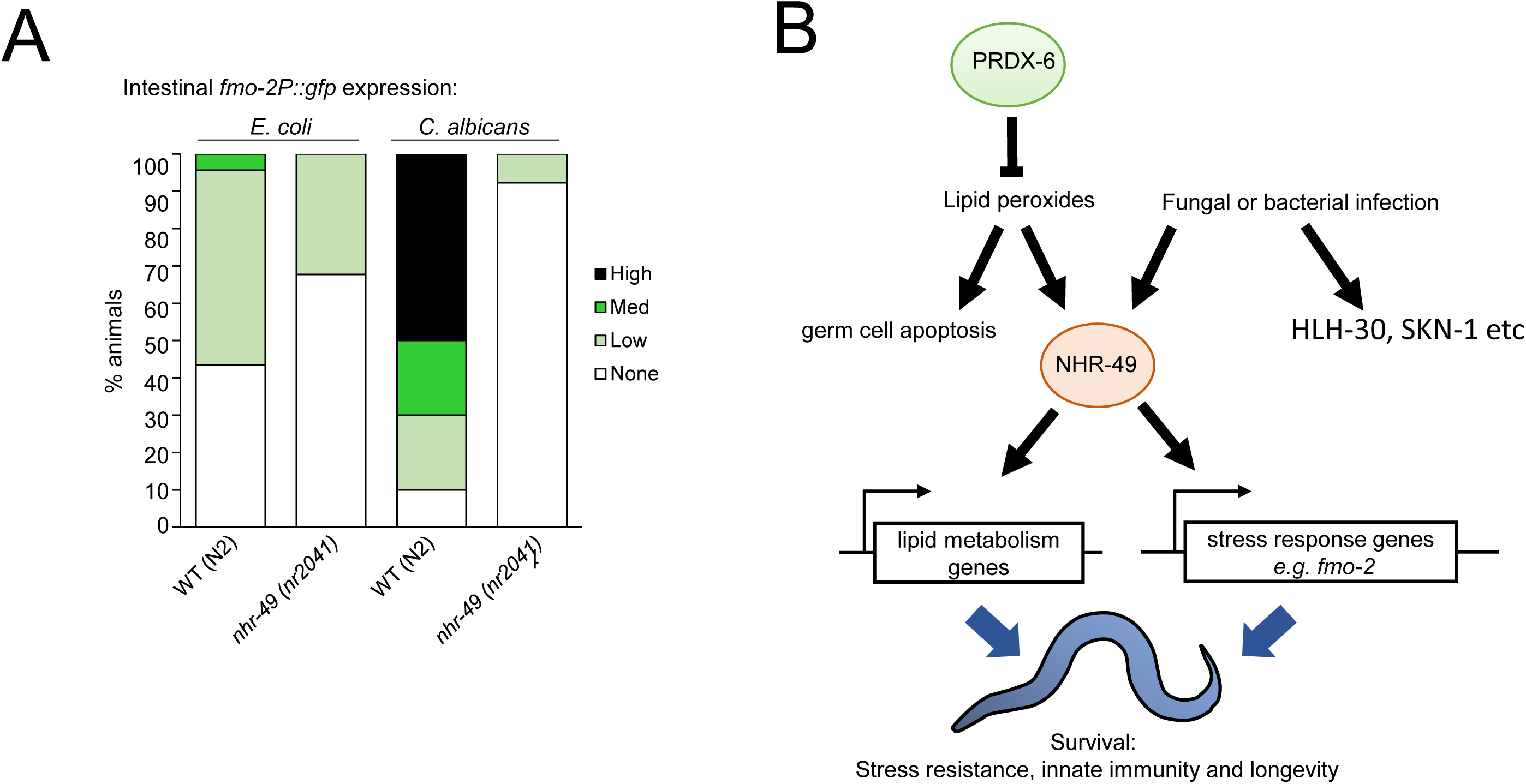
NHR-49 is required for induction of *fmo-2* in response to *C. albicans* leading us to propose that NHR-49 mediates the effect of loss of *prdx-6* on *fmo-2* expression an stress resistance and innate immunity. **[A]** The intestinal expression of the *fmo-2P::gfp* reporter gene was increased in response to 18h *C. albicans* exposure in wild-type but not *nhr-49* (*nr2041*) mutant *C. elegans* The experiment shown is representative of multiple independent experiments involving at least 20 animals in each group. There was a significant difference between intestinal *fmo-2p::gfp* expression in wild-type and *nhr-49* mutant *C. elegans* exposed to *C. albicans* (Chi^2^ test; p= 3.10 x10^-7^).**[B]** We propose that loss of PRDX-6’s lipid peroxidase activity leads to increased lipid peroxides which may increase NHR-49 activity, activating a transcriptional program that includes the flavin monooxygenase, FMO-2, thus prolonging survival during acute oxidative stress, fungal and bacterial infection. Fungal and bacterial infection also activate other transcriptional regulators (HLH-30, SKN-1 etc) that contribute to innate immunity.

## Discussion

Here we have uncovered that the single *C. elegans* 1-Cys Prdx, PRDX-6, plays important and unexpected roles in regulating oxidative stress resistance and innate immunity. Notably, we reveal that loss of this thiol peroxidase not only increases the lifespan of *C. elegans* but also prolongs survival following exposure to acutely toxic oxidative stress agents and two very different opportunistic human pathogens. To our knowledge, this is the first study to identify a role for a Prdx in suppressing innate immune defences and promoting survival under physiological and acute stress conditions.

Studies with Prdx6 knockout mice have suggested Prdx6 provides a non-redundant barrier to ROS and is important in maintaining normal insulin signalling (Pacifici *et al*, 2014), fertility (Ozkosem *et al*, 2015) brain (Phasuk *et al*, 2021a; Phasuk *et al*, 2021b) and muscle function (Pacifici *et al*, 2020). Consistent with this we have shown that *prdx-6* mutant *C. elegans* have increased ROS levels. Although, we found no evidence that this resulted in reduced insulin-signaling or affected feeding behaviour, our data are consistent with other work establishing that small increases in ROS can increase stress resistance and longevity (Schaar *et al*, 2015; Schulz *et al*, 2007; Scialo *et al*, 2020; Yang & Hekimi, 2010; Zarse *et al*, 2012). In this case, our data suggest that this increased survival under stress conditions may require the transcriptional regulator NHR-49. This adds to the evidence that this HNF4/PPARα-related nuclear hormone receptor’s role regulating lipid metabolism plays an important role in enhancing survival under various stress conditions (Doering *et al*, 2023).

Conserved immune pathways, such as the PMK-1 pathway, are important for the innate immunity of *C. elegans* to all pathogens (Aballay *et al*, 2003; Bolz *et al*, 2010; Pukkila-Worley *et al*., 2011; Shivers *et al*, 2010; Sifri *et al*, 2003; Troemel *et al*, 2006). However, transcriptomic data generated from worms infected with different pathogens show limited overlap in the genes that are up- and down-regulated (Irazoqui *et al*, 2010; Pukkila-Worley *et al*., 2011; Wong *et al*, 2007; Dasgupta *et al*, 2020; Wani *et al,* 2021). Thus, although there are some core immune pathways, unique responses are also generated in response to different infectious agents. Here we show that the Flavin monoxygenase, FMO-2, identified as a component of the NHR-49 up-regulated defence during *S. aureus, E. faecalis* and *C. neoformans* infection (Dasgupta *et al*, 2020; Wani *et al*., 2021), also represents a shared component of the innate immune defence to the fungal pathogen *C. albicans*. Intriguingly, although overexpression of *fmo-2* increased the survival of *C. elegans* infected with *C. albicans*, wild type and *fmo-2* mutant *C. elegans* that survived the first day succumbed to infection at similar rates. Similarly, although NHR-49 is important, loss of *fmo-2* alone does not increase the susceptibility of *C. elegans* to the gram negative bacteria *Pseudomonas aeruginosa* (Naim *et al*, 2021). Thus, a combination of upregulated genes most likely protect *prdx-6* mutant *C. elegans* against *C. albicans*. Similarly, it is likely that other genes upregulated in *prdx-6* mutant animals contribute to their increased stress resistance and extended lifespan.

The mechanism by which FMO-2 protects against infection is unclear. Notably, the mechanism by which this Flavin monoxygenase mediates the life-extending effects associated with dietary restriction and hypoxia also remain to be established (Huang et al, 2021). Although increased *fmo-2* extends the lifespan of *C. elegans* maintained on dead bacteria (Leiser *et al*., 2015), our data are consistent with other work suggesting that levels of this Flavin monoxygenase also affects interactions with the microbiome (Dasgupta *et al*., 2020; Wani *et al*., 2021). Consistent with the increased oxidative stress resistance of *prdx-6* mutant animals, high levels of FMO-2 protect against oxidative stress, potentially via increased stress-induced activation of the JNK pathway that plays important roles in stress resistance, apoptosis and innate immunity (Huang *et al*., 2021; Fabian *et al*, 2021).

NHR-49 levels are regulated by transcriptional and post-transcriptional mechanisms (Goh *et al*., 2018; Ratnappan *et al*., 2014). Consistent with previous studies we have not observed any increase in NHR-49 levels in response to infection (Wani *et al*., 2021) or in *prdx-6* mutant *C. elegans*. However, NHR-49 activity may also be increased by increasing levels of its natural ligand, which is likely to be an oxidised lipid (Doering *et al*., 2023). The phospholipid peroxidase activity of Prdx6 has been shown to make an important contribution to the metabolic reduction of lipid peroxides (Fisher *et al*., 2018). Moreover, loss of Prdx6 has been found to affect the synthesis of lipid-based signaling molecules in mammals (Kuda *et al*, 2018). Hence, it is possible that increased NHR-49 activity/*fmo-2* expression in *prdx-6* mutant *C. elegans* could reflect an increase in NHR-49’s natural ligand. In which case, future work to investigate the effect of *prdx-6* on *C. elegans* lipidome may provide important insight into the ligand-based activation of NHR-49.

Recent work has suggested that the main function of the redox-active cysteine in mammalian Prdx6 could be to mediate incorporation of selenium into selenoproteins, such as glutathione peroxidases (Fujita *et al*., 2024). The *C. elegans* genome encodes a single, selenoprotein, thioredoxin reductase, loss of which does not increase lifespan or innate immunity (Stenvall *et al*, 2011). From this we conclude that the phenotypes we describe for the *prdx-6* mutant are unlikely to reflect any effect on selenocysteine metabolism. Instead, our study provides evidence that the thiol peroxidase activity of a 1-Cys Prdx can provide an important barrier to ROS accumulation. As such, Prdx6 also limits expression of at least one gene associated with increased lifespan and innate immunity. Significantly, this highlights the important positive roles that low levels of ROS can play in maintaining health.

## Materials and Methods

### *C. elegans* strains

The following strains were used in this study: wild-type N2; VE2 *prdx-2* (*gk169*) II; VC1668 *fmo-2* (*ok2147*) IV; VE40 *N2 eavEx20[fmo-2p::gfp + rol-6(su1006)]*; VE43 *nhr-49* (*nr2041*) *eavEx20[fmo-2p::gfp + rol-6(su1006)]*; LX960 *lin-15B&lin-15A(n765) X;vsIs97 [tph-1p::DsRed2 + lin-15(+)]*. *C. elegans* bearing *prdx-6* (*tm4225*) IV and *prdx-6* (*tm4284*) IV were obtained from the Mitani lab and outcrossed 6x with our wild-type N2 strain to generate VE44 *prdx-6* (*tm4225*) IV and VE45 *prdx-6* (*tm4284*) IV. VE40 and VE44 were outcrossed to generate VE46 *prdx-6(tm4225) eavEx20[(fmo-2p::gfp + rol-6(su1006)].* VE48 *N2 eavEx21[prdx-6p::gfp + rol-6(su1006)]* was generated by microinjection of wildtype (N2) with pPD95.67+prdx-6P::GFP (containing the 2.5kb immediately upstream of the N-terminus of *prdx-6* plus the first 22 amino acids of *prdx-6* cloned in frame with GFP) together with the dominant co-injection marker *rol-6* (*su1006*). pPD95_67 was a gift from Andrew Fire (Addgene plasmid # 1490; http://n2t.net/addgene:1490; RRID:Addgene_1490). VE49 *N2 eavEx21[prdx-6p::gfp + rol-6(su1006)] vsIs97 [tph-1p::DsRed2 + lin-15(+)]* was generated by crossing VE48 and LX960. VE40, VE43, VE46, VE48 and VE49 were maintained by selection of worms with the roller phenotype. MD701 bcIs39 [lim-7p::ced-1::GFP + lin-15(+)] V was crossed with VE44 to generate VE50 *prdx-6* (*tm4225*) IV bcIs39 [lim-7p::ced-1::GFP + lin-15(+)] V or with VE45 to generate VE51 *prdx-6* (*tm4284*) IV bcIs39 [lim-7p::ced-1::GFP + lin-15(+)] V.

### Imaging and analysis of animals expressing fluorescent protein-encoding transgenes by fluorescent microscopy

Animals were transferred into M9 buffer containing 0.06% levamisole (Sigma L9756) for immobilization on 2%–2.5% (w/v) agarose pads. Fluorescence was observed using an epifluorescent Zeiss Axioskop 2 compound microscope and imaged under appropriate filters using AxioVision software (version 3.1.2.1). Intestinal expression of *fmo-2p::gfp* was assessed based on the number of GFP-positive intestinal nuclei as high(>12), medium (6-11), low (1-6) or none(no expression) as described previously (Crook-McMahon *et al*, 2014). CED-1::GFP engulfed corpses within the most visible arm of the gonad were counted.

### Detection of reactive oxygen species by H_2_DCFDA staining

Approximately 500 synchronised young adult *C. elegans* were washed off a NGM-L plate in 1ml of M9 buffer in to a microfuge tube. Animals were allowed to settle for approximately 3 min to form a pellet before the supernatant was removed. Animals were washed with 1ml of M9 buffer for 5 min two further times to remove any bacteria then incubated in 250μl of 25μM H_2_DCFDA diluted in M9 buffer, on a gently shaking platform in the dark for 30 min. Worms were allowed to settle and the supernatant removed. Worms were washed for 5 min with M9 buffer, the supernatant removed, and then a 30 min wash in M9 buffer followed by a final 5 min wash in M9 buffer. The supernatant was then removed and worms were gently resuspended in 20μl of 0.06% levamisole in M9 before mounting on agarose pads and imaging using a Zeiss Axioskop 2 fluorescent microscope.

### Analysis of lifespan

Approximately 150 L1 larval stage worms were allowed to develop to L4 larval stage at the described temperature before being transferred on to four separate plates. Worms maintained at 25°C were moved to fresh plates daily, or on alternate days if at 15°C or 20°C, until progeny had finished being laid. Survival was observed on the indicated days with worms not responding to prodding with a platinum wire scored as dead and removed from the plate.

### Analysis of oxidative stress resistance

30 L4 larval stage worms were transferred to freshly prepared NGM-L plates containing 5mM hydrogen peroxide (Sigma) or 10mM sodium arsenite along with a small amount of *E.coli* (OP50). Survival was observed at the indicated time points with worms not responding to prodding with a platinum wire scored as dead and removed from the plate.

### Infection experiments

Synchronised L4/Young adult animals were transfered on to an unseeded NGM-L plate for 1-1.5h before being transferred on to plates seeded with the pathogen. Survival was then monitored daily at 25°C. For *S*. Typhimurium experiments *cdc25.1* RNAi-treated L4/young adult animals were used. *Staphylococcus aureus*-seeded plates were prepared essentially as described previously Sifri et al 2003. A single colony of *S. aureus* (NCTC 8325) was inoculated in 10ml of Tryptic Soy liquid media (Fluka) supplemented with 5ug/ml Nalidixic acid (Sigma-Aldrich) and grown overnight at 37°C on a shaking platform. 10μl of the *S. aureus* (NCTC8325) culture was spotted onto TS supplemented with 5ug/ml Nalidixic acid agar plates and plates were dried overnight at 37°C. *Candida albicans*-infection experiments were carried out essentially as described by (Pukkila-Worley *et al*., 2011). A single colony of *C. albicans* (SN148) was inoculated in 10ml of Yeast Peptone Dextrose (YPD) liquid media, and grown overnight at 30°C on a rotating wheel then 10μl of this culture was seeded onto BHI media (BBL (6.0g/L Brain Heart, Infusion from (solids), 6.0g/L Peptic digest of Animal Tissue, 5.0g/L Sodium Chloride. 3.0g/L Dextrose, 14.5g/L Pancreatic Digest of Gelatin, 2.5g/L Disodium Phosphate) and 10g/L Bacto Agar (BD) supplemented with 45ug/ml kanamycin (Sigma-Aldrich). For survival assays, *S. aureus*-seeded TS plates and *C. albicans*-seeded BHI plates were spotted with 8-16μl of 40mM 5’Fluoro-2’-deoxyuridine (FUDR) immediately prior to addition of *C. elegans.* Survival was observed at the indicated time points with worms not responding to prodding with a platinum wire scored as dead and removed from the plate.

### Statistical analysis

Experimental data shown in each figure panel are representative of at least three independent biological repeats. Log Rank analysis was used to identify any statistically significant differences in survival data. Chi^2^ tests were used to determine the significance of differences between the intestinal expression profiles for transcriptional reporters e.g. *fmo-2*::*gfp* under different conditions or in different genetic backgrounds. Student’s T tests were used to analyse differences between the number of ced-1::GFP engulfed nuclei in different groups.

## CRediT author statement

**Emma Button:** Conceptualisation, Investigation, Visualisation, Methodology, Formal analysis, Writing -original draft. **Jake Lewis:** Investigation, Formal analysis, Validation. **Emilia Dwyer:** Methodology, Investigation, Validation, Writing-review and editing. **Elise McDonald**: Investigation, Validation. **Eloise Butler:** Investigation. **Fiona Pearson**: Investigation. **Elizabeth Veal:** Conceptualisation, Supervision, Funding acquisition, Visualisation, Methodology, Project administration, Writing original draft, review and editing.

## Acknowledgements

We are grateful to the National Bioresource Project for the nematode c/o Dr. Shohei Mitani for generating animals bearing the *prdx-6* mutant alleles used in this study. Some strains were provided by the CGC, which is funded by NIH Office of Research Infrastructure Programs (P40 OD010440). Prof. Janet Quinn and Dr Yasmin Ahmed kindly provided the *Candida albicans* and Dr. Judith Hall provided the *Staphylococcus aureus* used in this study. We are very grateful to Antonio Miranda Vizuete and Janet Quinn for comments on this manuscript. This work was supported by an MRC-funded studentship (EB), MRC grant G0800082 (EV), BB/T002484/1 (EM), BB/T008695/1 (ED) and Newcastle University.

