## Supplementary Figures and Tables for "The 1 –Cys peroxiredoxin, PRDX-6, suppresses a pro-survival response, including the Flavin monoxygenase, FMO-2, that protects against fungal and bacterial infection"

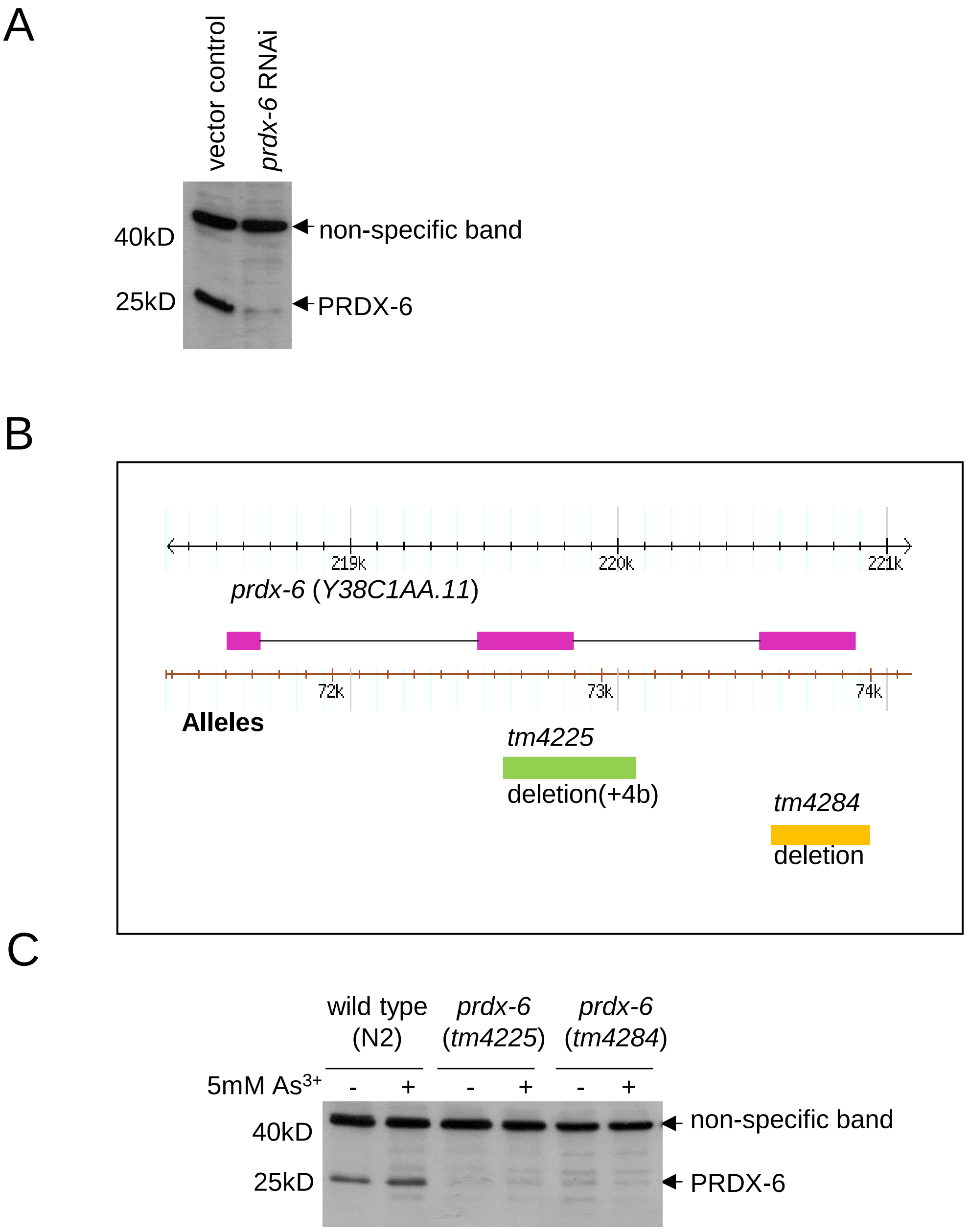


**Fig. S1 Antibody 5754-1-4M234/4J10_120208 detects a protein predicted to be PRDX-6 that is reduced by *prdx-6* RNAi and absent from *prdx-6* mutant worms.** [A] Immunoblot of wild type worms treated with vector control and *prdx-6* RNAi with anti-PRDX-6 antibodies indicates that *prdx-6* RNAi reduces the levels of a protein of the expected size(25.6kDa). [B] The genome structure of the *prdx-6* gene indicating the regions that are deleted in worms bearing the *tm4225* and *tm4284* alleles [C] Western blot of protein lysates of wild type, (N2) *prdx-6 (tm4225)* and *prdx-6 (tm4284)* mutant worms before and after treatment with 5mM arsenite for 5 minutes and probed with antibody 5754-1-4M234/4J10_120208. The absence of the 25kDa band from lysates from *prdx-6 tm4225* and *tm4284* mutants confirming loss of the PRDX-6 protein. A non-specific band around 40kD is also indicated, confirming even protein loading.

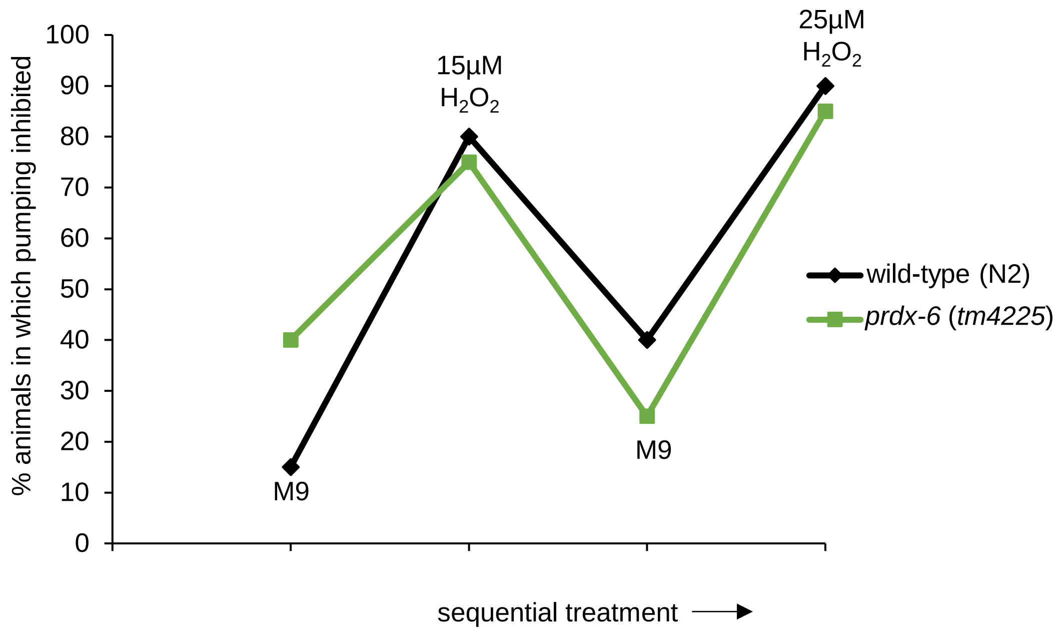


**Fig. S2 PRDX-6 is not required for inhibition of pumping in response to $H_2O_2$.** When young adult animals were sequentially exposed to a droplet of M9 (control), 15µM $H_2O_2$, M9 then 25µM $H_2O_2$ a similar % of wild-type and *prdx-6* (*tm4225*) mutant animals exhibited transient inhibition of pumping in response to $H_2O_2$ (n=20 in each group). Pumping inhibition was assayed essentially as previously described (Bhatla and Horvitz, 2015): A droplet of M9 or M9 containing hydrogen peroxide was added close to the animal, using a pipette or a needle, such that the liquid engulfed the head only. The response was scored by eye on a stereoscope and inhibition was determined upon a noticeable pause in the rhythmic pumping of the pharyngeal grinder within 10s of liquid entering the pharynx. Experiment was repeated with similar results and a representative experiment is shown.

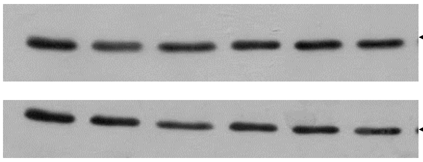


**Fig. S3 Wild type and *prdx-6* mutant worms contain similar levels of PRDX-2.** Western blot analysis of wildtype, *prdx-6* (*tm4225*) and *prdx-6* (*tm4284*) mutant worms before and after 5 min exposure to 5 mM $H_2O_2$ with anti-PRDX-2 antibodies (Olahova *et al*, 2008). The tubulin antibody was used as a loading control to normalise PRDX-2 levels using ImageJ software. The average quantified levels of PRDX-2 compared to the unstressed wild type from two repeats are displayed beneath the Western Blot. Student's T-test revealed that none of the groups were significantly different to wild type unstressed, $p<0.05$

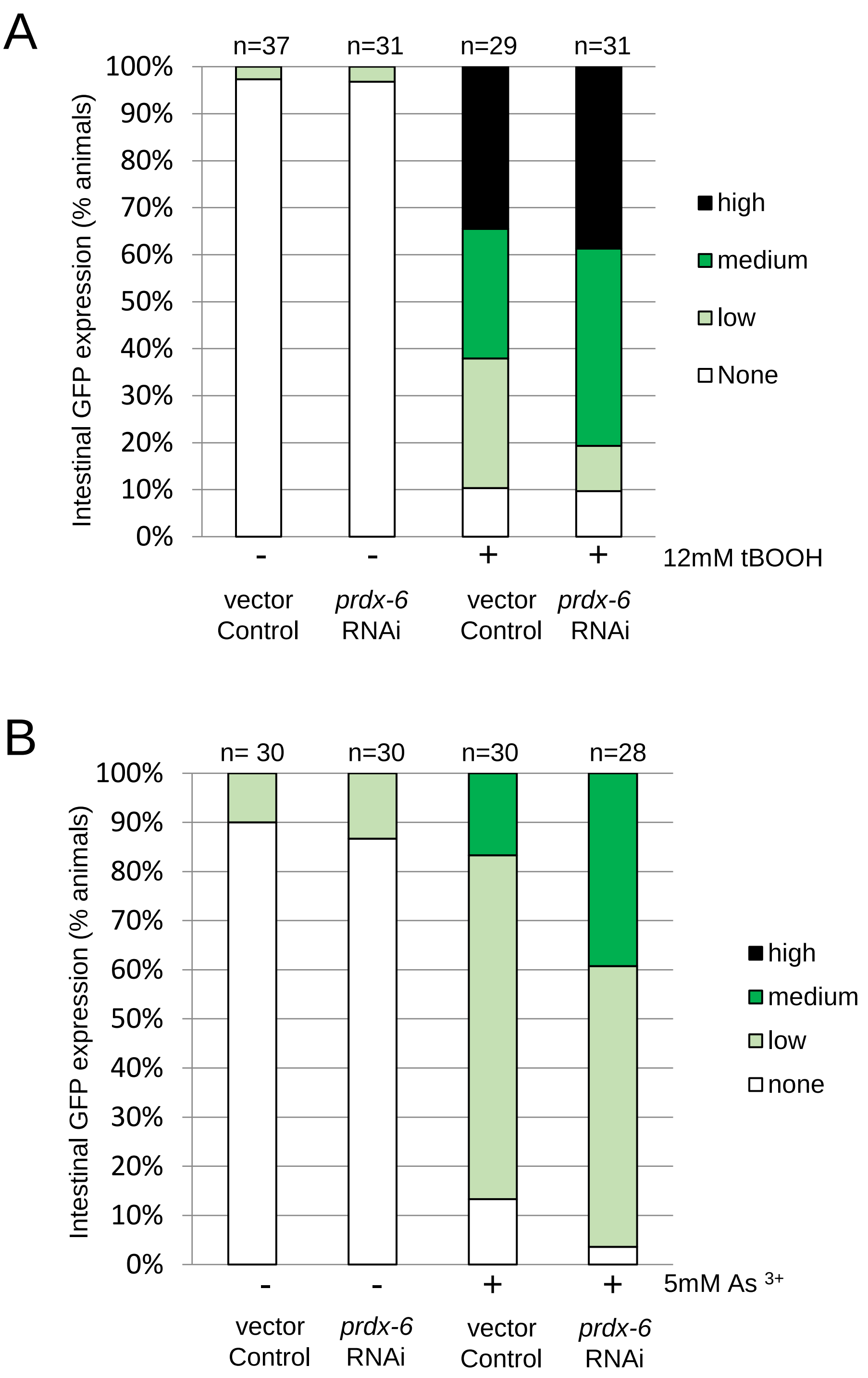


**Fig. S4 *prdx-6* RNAi does not significantly effect basal, tBOOH- or arsenite- induced intestinal *gcs-1*P::GFP expression.** The intestinal expression of *gcs-1*P::GFP was scored in L4 wild type animals treated with either vector control or *prdx-6* RNAi, without stress and stressed with [A] 12mM tBOOH or [B] 5mM arsenite ($As^{3+}$) The experiment was performed twice with similar results and a representative experiment is shown. Chi-squared test compared vector control treated worms to *prdx-6* RNAi treated, [A] without stress p=0.90; with tBOOH stress p=0.31 [B] without stress p=0.69; with arsenite stress p=0.097 n= number of animals per group.

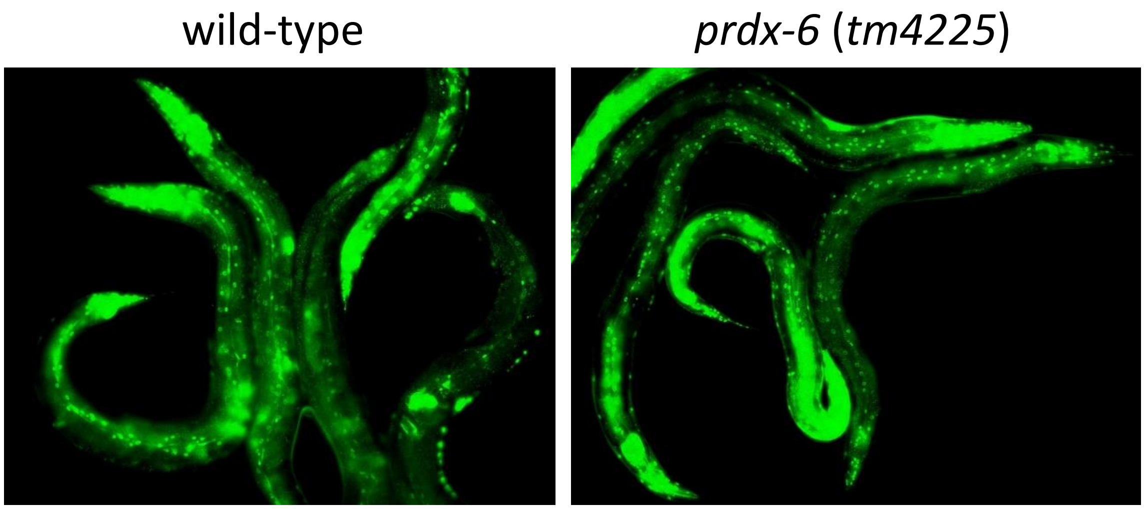


**Fig. S5 NHR-49+GFP levels and distribution are similar in *wild type and prdx-6 mutant C. elegans*** NHR-49::GFP expression was observed in well-fed wild-type (AGP24) and *prdx-6* (*tm4225*) mutant animals expressing *Pnhr-49::nhr-49::gfp* (Ratnappan et al. 2014). More than 100 adult animals were examined and imaged under identical conditions/exposures. Although there was some variation between animals in each group, no differences were observed.. Representative animals from each group are shown.

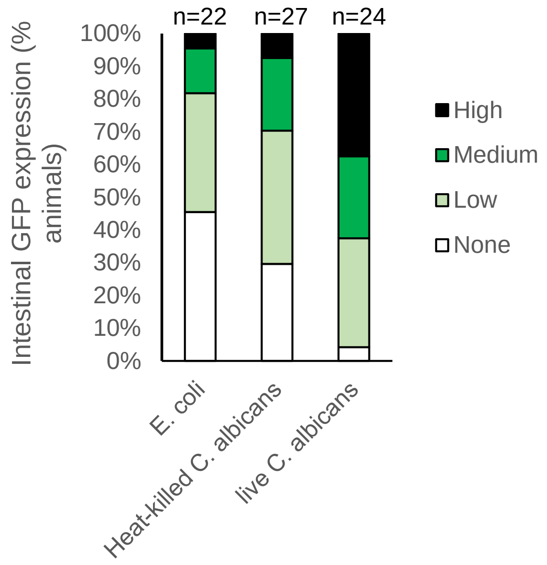


**Fig. S6 *fmo-2P::gfp* expression in wild type worms on *E. coli* or following 6h exposure to live or heat-killed *C. albicans.*** The intestinal expression of *fmo-2P*::GFP was scored in L4 wild type animals maintained on *E. coli* (OP50) 6hour following transfer to *E.coli,* live or heat-killed *C. albicans.* $Chi^2$ test comparing expression in animals treated with E. coli or heat-killed *C. albicans* p=0.674 whereas exposure to live *C. albicans* significantly increased intestinal *fmo-2P*::GFP expression compared with *E. coli* ($chi^2$ test p=0.0021) or heat-killed *C. albicans* ($chi^2$ test p=0.017). n=number of animals per group.

**Table S1** Lifespan parameters associated with survival monitoring experiment shown in Fig. 4C (lifespan at 15°C)

| **Strain** | **Mean** (days) | **Median** (days) | **Log Rank p-value (relative to wildtype)** |
|---|---|---|---|
| wild type (N2) | 24.12 | 24 | |
| *prdx-6* (*tm4225*) | 30.63 | 32 | <0.001 |
| *prdx-6* (*tm4284*) | 33.45 | 32 | <0.001 |

**Table S2** Lifespan parameters associated with survival monitoring experiment shown in Fig. 4D (lifespan at 25°C)

| **Strain** | **Mean** (days) | **Median** (days) | **Log Rank p-value (relative to wild type)** |
|---|---|---|---|
| wild type (N2) | 6.69 | 6 | |
| *prdx-6* (*tm4225*) | 7.34 | 7 | 0.447 |
| *prdx-6* (*tm4284*) | 7.61 | 7 | 0.132 |

**Table S3** Combined lifespan parameters from 3 independent lifespan experiments at 15°C and 25°C

| | **15°C** | | | **25°C** | | |
|---|---|---|---|---|---|---|
| **Strain** | Mean Lifespan (days) | Median Livespan (days) | no. of worms | Mean Lifespan (days) | Median Livespan (days) | No. of worms |
| N2 | 24.7 | 24 | 187 | 7.26 | 7 | 112 |
| *prdx-6* (*tm4225*) | 29.1 | 30 | 159 | 7.50 | 7 | 115 |
| *prdx-6* (*tm4284*) | 29.5 | 30 | 156 | 7.40 | 7 | 108 |

**Table S4** Loss of PRDX-6 causes an increased sensitivity to *S*. Typhimurium (parameters related to experiment shown in Fig. 5A)

| **Strain** | **no. of worms** | **Mean survival** (days) | **Log Rank p-value (relative to wild type)** |
|---|---|---|---|
| wild type (N2) | 75 | 6.87 | |
| *prdx-6* (*tm4225*) | 57 | 5.07 | <0.001 |
| *prdx-6* (*tm4284*) | 69 | 5.17 | 0.001 |

**Table S5** Loss of PRDX-6 causes an increased resistance to *S. aureus* (parameters related to experiment shown in Fig. 5B)

| **Strain** | **no. of worms** | **Mean survival** (days) | **Log Rank p-value (relative to wild type)** |
|---|---|---|---|
| wild type (N2) | 70 | 1.77 | |
| *prdx-6* (*tm4225*) | 66 | 3.47 | <0.001 |
| *prdx-6* (*tm4284*) | 63 | 3.03 | <0.001 |

**Table S6** Loss of PRDX-6 causes an increased resistance to *C. albicans* (parameters related to experiment shown in Fig. 5C)

| **Strain** | **no. of worms** | **Mean survival** (days) | **Log Rank p-value (relative to wild type)** |
|---|---|---|---|
| wild type (N2) | 109 | 2.55 | |
| *prdx-6* (*tm4225*) | 125 | 3.86 | <0.001 |
| *prdx-6* (*tm4284*) | 112 | 3.55 | <0.001 |

**Table S7** FMO-2 increases resistance to *S. aureus* (parameters related to experiment shown in Fig. 6D)

| **Strain** | **no. of worms** | **Mean survival** (days) | **Log Rank p-value (relative to wild type)** |
|---|---|---|---|
| wild type (N2) | 113 | 2.86 | |
| *fmo-2* (*ok2147*) | 112 | 2.22 | <0.001 |
| *fmo-2oEx* | 116 | 3.22 | <0.001 |

**Table S8** FMO-2 increases resistance to *C. albicans* (parameters related to experiment shown in Fig. 6E)

| **Strain** | **no. of worms** | **Mean survival** (days) | **Log Rank p-value (relative to wild type)** |
|---|---|---|---|
| wild type (N2) | 98 | 2.32 | |
| *fmo-2* (*ok2147*) | 89 | 1.91 | =0.008 |
| *fmo-2oEx* | 92 | 4.09 | <0.001 |
